# Methyl-substituted 1,10-phenanthroline derivatives in copper(II) complexes enhance antitumor activity and inhibit NHE1 in MDA-MB-231 breast cancer cells

**DOI:** 10.64898/2026.09.04.749506

**Authors:** K.S. Muñoz Garzón, V. de Giusti, C.Y. Fernandez, G. Facchin, V. Martínez, A.L. Di Virgilio

## Abstract

Three copper(II) complexes containing 1,10-phenanthroline derivatives, [CuCl₂(phen)].0.5H₂O (**1**), [CuCl₂(neo)].0.75H₂O (**2**), and [CuCl₂(tmp)].H₂O (**3**), were evaluated for their antitumor activity in MDA-MB-231 breast cancer cells. All complexes markedly reduced cell viability, exhibiting significantly lower IC₅₀ values and higher selectivity indices than the corresponding free ligands, CuCl₂, and cisplatin, highlighting the therapeutic advantage of metal complexation. Among them, only **1** induced DNA damage at sub-IC₅₀ concentrations.

Cytotoxicity in all complexes was associated with intracellular ROS generation and apoptosis, although **2** and **3** produced a stronger oxidative response. Complex **3** additionally promoted necrosis. Cell proliferation was inhibited in a concentration-dependent manner, accompanied by increased intracellular copper accumulation.

Moreover, **2** and **3** significantly inhibited Na⁺/H⁺ exchanger 1 (NHE1) activity, reducing cell migration and MMP-9 activity. Western blot analysis further demonstrated that all three complexes modulated the expression of NHE1, G protein-coupled estrogen receptor (GPER), and apoptosis-related proteins.

Overall, our findings show that ligand methylation enhances the antitumor activity of copper(II) complexes and shifts their mechanism of action from DNA damage-driven cytotoxicity toward ROS-mediated apoptosis, while also improving inhibition of NHE1-dependent migratory pathways. These results identify NHE1 as a key molecular target underlying the enhanced antitumor activity of methylated phenanthroline copper(II) complexes.

## Introduction

Breast cancer is the most commonly diagnosed cancer and the second leading cause of cancer-related mortality worldwide in 2024 [1]. Among its molecular subtypes, triple-negative breast cancer (TNBC) is characterized by the absence of estrogen receptor, progesterone receptor, and human epidermal growth factor receptor 2 (HER2) expression. TNBC accounts for approximately 15–20% of breast cancer cases and is associated with an aggressive clinical course, early metastatic dissemination, and poor prognosis [2]. Unlike hormone receptor-positive breast cancer, TNBC lacks effective targeted therapies, leaving chemotherapy, immunotherapy, antibody-drug conjugates, and poly(ADP-ribose) polymerase inhibitors as the main systemic treatments. However, intrinsic and acquired drug resistance often compromises efficacy, underscoring the need for novel therapeutic approaches [3].

Over the past decades, researchers have made considerable efforts to develop metal-based anticancer agents, particularly copper(II) complexes, which have shown remarkable antitumor activity *in vitro* and *in vivo.* Copper complexes containing nitrogen-donor ligands, including 1,10-phenanthroline derivatives, have shown promising cytotoxic effects against different cancer cell types, supporting their potential as candidates for anticancer drug development [4–6]. Many retain activity against cisplatin-resistant tumor cells and can even eliminate cancer stem cells [7–9]. Importantly, the biological activity of copper complexes can be strongly influenced by ligand structure, suggesting that structural modifications may modulate their cellular targets and mechanisms of action.

Beyond their cytotoxic activity, increasing evidence indicates that copper complexes can modulate signaling pathways involved in tumor progression. Among these pathways, the Na⁺/H⁺ exchanger isoform 1 (NHE1) is a key regulator of intracellular pH homeostasis and has emerged as an attractive therapeutic target in breast cancer [10,11]. NHE1-mediated proton extrusion promotes intracellular alkalinization while acidifying the extracellular microenvironment, thereby facilitating extracellular matrix remodeling, invasion, and metastasis [10,12]. Consistently, pharmacological inhibition of NHE1 suppresses breast cancer cell proliferation and migration while enhancing apoptosis. These findings support NHE1 inhibition as a potential strategy for limiting multiple processes associated with breast cancer progression [12,13]. Moreover, extracellular matrix (EDM) remodeling is largely mediated, in part, by matrix metalloproteinases (MMPs), including the gelatinases MMP-2 and MMP-9, which are associated with the invasive phenotype of breast cancer cells. NHE1 activity has been linked to ECM degradation and tumor cell invasion through invadopodial formation and matrix-degrading activity, while its inhibition reduces ECM degradation and invasion, supporting a functional association between NHE1 activity and extracellular protease-mediated tumor progression [14–17]

In addition to classical estrogen receptors, the G protein-coupled estrogen receptor (GPER) has emerged as an important mediator of rapid, non-genomic estrogen signaling in breast cancer. GPER is expressed in several breast cancer subtypes, including triple-negative breast cancer, and its activation has been associated with signaling pathways involved in cell proliferation, migration, invasion, and tumor progression, including EGFR/ERK signaling [18]. Interestingly, GPER interacts with Na+/H+ exchanger regulatory factor 1 (NHERF1), a scaffolding protein involved in regulating ion transport proteins and signaling pathways in breast cancer cells [19]. These findings raise the possibility that therapeutic strategies targeting breast cancer progression may modulate GPER expression, potentially contributing to their antitumor effects.

In our previous work, we demonstrated that copper(II) complexes containing phenanthroline derivatives exert potent antitumor activity in luminal breast cancer cells through a multitarget mechanism involving apoptosis, inhibition of NHE1 activity, and modulation of MMPs. We also showed that subtle changes in ligand architecture significantly influenced their biological activity [14].

Building on these findings, the present study investigates whether the copper(II) complexes containing only phenanthroline derivatives as primary ligands: 1,10-phenanthroline [CuCl₂(phen)]·0.5H₂O (**1**), neocuproine [CuCl₂(neo)]·0.75H₂O (**2**), and tetramethyl-phenanthroline [CuCl₂(tmp)]·H₂O (**3**) (Fig. 1), retain their antitumor activity in the highly aggressive triple-negative breast cancer cell line MDA-MB-231. Furthermore, we examined how ligand structure influences their mechanism of action by evaluating oxidative stress, apoptosis, DNA damage, NHE1 activity, clonogenic capacity, cell migration, MMP activity, and the expression of proteins associated with tumor progression.

**Figure 1.**
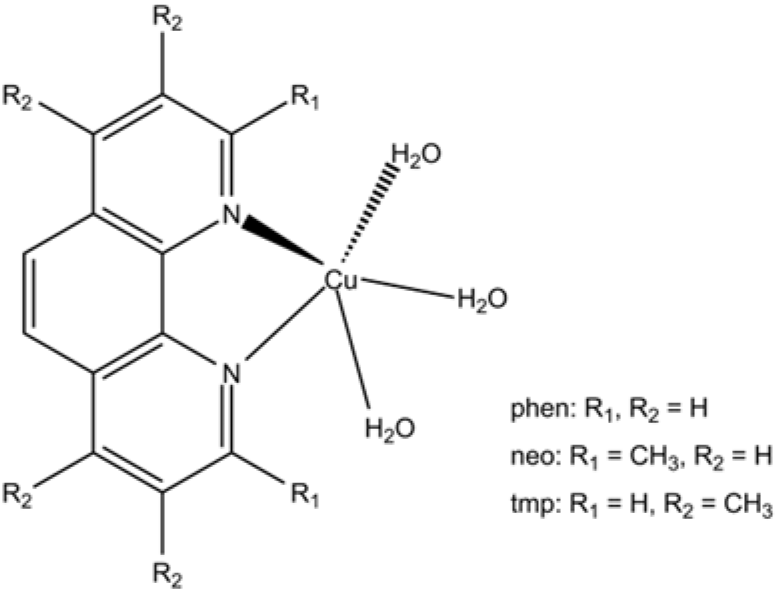
Scheme of the copper(II) complexes (as expected in solution).

## Experimental

### Materials

Tissue culture materials were purchased from Corning (Princeton, NJ, EUA) and APBiotech (Buenos Aires, Argentina). Dulbecco’s Modified Eagle Medium Nutrient Mixture F-12 (DMEM/F-12) and TrypLE™ were obtained from Gibco (Gaithersburg, MD, EUA) and fetal bovine serum (FBS) from Internegocios S.A. (Buenos Aires, Argentina); Rhodamine 123, SYBR Green, and low-melting-point agarose from Invitrogen Corporation (Buenos Aires, Argentina). Bleomycin (BLM) (Blocamycin) was kindly provided by Gador S.A. (Buenos Aires, Argentina). MDA-MB-231 and Hacat cell lines were purchased from ATCC®. Polyvinylidene fluoride (PVDF) membrane (BioRad, CA, USA), 2’,7’-bis-(2-carboxyethyl)-5-(and-6)-carboxyfluorescein acetoxymethyl ester (BCECF-AM) (ThermoFisher, USA), Xanthine, EDTA, Carbonate Buffer, Xanthine oxidase, NAC (N-Acetyl-L-Cysteine), gelatine B and nitroblue tetrazolium (NBT) (Sigma).

### Copper (II) Complexes

Three copper (II) complexes [CuCl_2_(phen)]·0.5H_2_O (**1**), [CuCl_2_(neo)]·0.75 H_2_O (**2**), [CuCl2(tmp)]·H_2_O (**3**), where phen stands for 1,10-phenanthroline, neo for neocuproine (2,9-dimethyl-1,10-phenanthroline) and tmp for 3,4,7,8-tetramethyl-1,10-phenanthroline, were previously synthesized and physiochemically characterized at the Faculty of Chemistry, University of the Republic, Montevideo, Uruguay, as previously reported [20–22].

## Methods

### Experimental elemental analysis for complex

To assess compound purity, we performed infrared (IR) spectroscopy and elemental analysis. IR spectra were recorded on a Shimadzu IR Prestige 21 (4000 to 400 cm−1) using 1% KBr pellets. The spectra agreed well with previously reported data [22–24]. Elemental analysis of C, N, and H was performed in a Thermo Flash 2000 elemental analyzer. The calculated and experimental elemental compositions were as follows: [CuCl_2_(phen)]·0.5H2O: Calc./Found for CuCl_2_C_12_H_9_N_2O_ 0,5 %C 44.53/44.18, %N 8.66/8.36, %H2.80/2.92; [CuCl_2_(neo)]·0.75 H2O: Calc./Found CuCl_2_C_14_H_13_,5N_2_O 0.75 %C: 47.20/47.16 %N: 7.86/7.62 %H: 3.82/3.86; [CuCl_2_(tmp)]·H_2_O: Calc. for CuC_16_H_18_N_2_OCl_2_/Found: %C: 49.43/49.09, %N: 7.21/7.08, %H: 4.67/4.61.

The elemental analysis results were consistent with the proposed molecular compositions of the complexes. Based on previous studies by our group, the diimines ligands in complexes **1**-**3** are expected to remain coordinated to the copper center in solution, with water molecules completing the coordination sphere [24].

### Cell line MDA-MB-231

The human breast cancer cell line (MDA-MB-231) derived from pleural effusions of a breast cancer patient and obtained from the American Type Culture Collection (ATCC®), was cultured in DMEM/F-12 medium supplemented with 10% fetal bovine serum (FBS), 100U/ml penicillin, and 100 μg/mL streptomycin at 37°C with 5% CO_2_ in a humidified atmosphere. Cells were initially seeded in a 75 cm^2^ culture flask, and when they reached 80% confluence, they were detached using TrypLE^TM^. Experiments were carried out in multiwell culture plates where cells were allowed to attach and treated for each complex.

### Cytotoxicity assay

Cell viability and half-maximal inhibitory concentration (IC_50_) values were determined by the 3-(4,5-dimethylthiazol-2-yl)-2,5-diphenyltetrazolium bromide (MTT) method. Cells were seeded in 96-well culture plates at a density of 1.5×10^4^ cells per well. After 24 h, the cells were treated with **1**, **2**, **3,** and the corresponding ligands, at concentrations ranging from 0.5 to 200 μM. Next, the medium was aspirated from the wells, and the MTT reagent was incubated in DMEM/F-12 for 3 h. The formazan crystals were dissolved in DMSO (100 μl), and absorbance was measured using a multi-plate reader Multiskan FC, Thermo Scientific at an excitation wavelength of 570 nm. The IC_50_ was determined through nonlinear regression analysis employing PrismGraph Software.

### Genotoxicity studies

To detect DNA strand breaks, the comet assay was used. Electrophoresis at high pH results in comet-shaped structures. Briefly, 1.5×10^5^ cells were cultured in 12-well plates for 24 h. The monolayer was treated with the complexes at different concentrations for an additional 24 h. A negative control used cells without complex, and bleomycin (10 μg/mL) served as the positive control. Cells were washed with phosphate-buffered saline (PBS), detached using TrypLe^TM^ and centrifuged at 2000 rpm at 4°C. The pellet was resuspended in 75 μL of low-melting-point agarose and added to slides containing normal-melting-point agarose, left to solidify at 4°C for 15 min, and immersed in lysis solution. Electrophoresis was performed under alkaline conditions in a horizontal electrophoresis chamber for 30 min at 25 V and 4°C. Following electrophoresis, slides were washed with a neutralization buffer and stained with SYBR Green. DNA damage was visualized using a Nikon Eclipse E200 Epifluorescence microscope. The images were analyzed to determine the tail moment using CometScore 1.5 software.

### Determination of reactive oxygen species (ROS) production

Reactive oxygen species (ROS) levels were measured using dihydrorhodamine 123 (DHR 123), a fluorogenic dye that turns into fluorescent rhodamine 123 when oxidized by cellular free radicals. MDA-MB-231 cells were seeded in 24-well plates and allowed to reach approximately 100% confluence. Next, cells were treated with different concentrations of complexes for 24 h. The monolayer was washed with Hank’s Balanced Salt Solution (138 mM NaCl, 5 mM KCl, 0,4 mM MgSO_4_·7H_2_O, 1.2 mM CaCl_2_·H_2_O, 0.4 mM KH_2_PO_4_, 4.1 mM NaHCO_3,_ and 5.5 mM C₆H₁₂O₆), and next, 480 μL of DHR-123 was added to each well and incubated for 30 min at 37°C in the dark. Cells were washed with Hank’s solution and incubated for 45 min at 37°C with lysis solution (0,1% Triton X-100). DHR-123 oxidation was detected by spectrofluorometry. ROS levels were normalized to total protein content, measured by the bicinchoninic acid (BCA) method. This method determines protein content based on the reduction of Cu^2+^ to Cu^+^ by proteins in an alkaline medium. The resulting cuprous ions react with BCA to produce a deep violet color, which was measured at 570 nm.

To further assess the involvement of ROS in the effects induced by the complexes, the monolayer was pretreated with 250 μM NAC for 2 h, next the culture medium was replaced with concentrations of the complexes. In both cases, cytotoxicity was evaluated by the MTT method. In addition, the effects of the complexes on antioxidant defenses were evaluated by determining SOD activity, GSH levels, and catalase activity.

### GSH and GSSG assay

GSH was determined using a method based on [25]. The cells were seeded in 12 well plates and treated for 24 h with the complexes. The cells were washed with PBS and lysed for 5 min at room temperature with 0.1% Triton X-100. For GSH determination, 100 µL aliquots of lysed cells were mixed with 1.8 mL of ice-cold phosphate buffer (Na_2_HPO4 0.1 M-EDTA 0.005 M, pH 8) and 100 µL of ophthaldialdehyde (OPT) (0.1% in methanol) and were incubated for 15 min under light protection. For GSSG determination, 100 µL aliquots of cells were first incubated with 0.04 M of N-ethylmaleimide (NEM) for 20 min in the dark (to avoid GSH oxidation). Then, the aliquots were mixed with 1.7 µL of 0.1 M NaOH and 100 µL OPT in the dark and on ice. The fluorescence was determined (λ_ex_ 350 nm and λ_em_ 420 nm) in a spectrophotometer Varioskan LUX (Thermofisher). Protein content in each cellular extract was quantified using the Bradford assay [26].

### Catalase activity

The effects of the copper(II) complexes on catalase activity were evaluated by measuring H₂O₂ degradation in cell lysates. Briefly, 10 μL of cell lysate was mixed with 20 μL of H_2_O_2_ 10 mM and incubated for 2 min at 37°C. To quantify residual hydrogen peroxide, 20 µL of ammonium metavanadate (NH_4_VO_3_) was added and incubated for 10 min at room temperature. Residual hydrogen peroxide reacts with ammonium metavanadate, forming a red-orange peroxovanadium complex. Absorbance was measured at 452 nm using a Varioskan LUX spectrophotometer (Thermofisher). A standard and blank was prepared simultaneously [27].

### SOD activity

The superoxide dismutases (SODs) are key antioxidant enzymes that protect cells from oxidative stress by catalyzing the dismutation of superoxide radical (O₂•⁻) into H_2_O and O_2_. The major isoforms are found in the cytosolic (SOD1 or Cu-Zn-SOD) and mitochondrial (Mn-SOD SOD_2_) regions. SOD activity was evaluated by the inhibition of nitroblue tetrazolium (NBT) reduction by superoxide radicals generated through the xanthine/xanthine oxidase system. The mixture of solutions contained xanthine (0.1 mM, pH 10.2), EDTA (25 mM pH 8), carbonate buffer solution (50 mM at pH 10.2), and NBT (0.025 mM), which were mixed with different concentrations of cell lysates (Triton X-100 0.1%) obtained from cells previously treated with the three complexes at the indicated concentrations. Xanthine oxidase was added before measurement. Absorbance was recorded at 560 nm every 30 s for 5 min using a Varioskan LUX (Thermo Fisher Scientific) spectrophotometer.

### Quantification of intracellular Copper

MDA-MB-231 cells were seeded until reaching a 80-90% confluent monolayer and treated with the copper complexes at the IC_50_ for 24 h. After incubation, the supernatant medium was transferred to 15 mL plastic centrifuge tubes, and the cell suspension was centrifuged for 10 minutes at 500 rpm and the supernatant was discarded. The cells were detached with TryplE^TM^ and transferred to the labeled centrifuge tubes. The samples were centrifuged (500 rpm for 10 min), and the supernatant was discarded. To remove extracellular copper, the cells were washed twice with 10 mL of cold PBS and centrifuged at 500 rpm for 10 min. Finally, the cells were resuspended in 2 mL of PBS and centrifuged again under the same conditions. The supernatant was carefully discarded. The cell pellets were treated with 150 μL of a 0.1% (v/v) Triton X-100 solution. The cell suspension was vortexed for 10 minutes to induce cell lysis, and a 10 μL aliquot was separated to determine the protein concentration. Next, 1 mL of 20% nitric acid solution was added to each sample. The mixtures were vortexed, and sonicated at maximum power for 60 minutes at 55°C in an ultrasonic bath, followed by refrigeration for 48 h. Samples were then sonicated again at 55°C for 2 h and immediately centrifuged at 3000 rpm for 20 min. 1000 μL of each sample were transferred to new 15 mL centrifuge tubes and analyzed using a Shimadzu ICPE-9820 inductively coupled plasma atomic emission spectrometer (PlaPiMu LaSeISiC Unit (CIC–UNLP)).

### Acridine orange/propidium iodide staining

Cellular and nuclear morphological alterations were evaluated using acridine orange (AO) and propidium iodide (PI) staining. MBA-MB-231 cells were grown in 12-well plates and incubated overnight. Copper(II) complexes were added at their respective IC_50_ for 24 h. Cells were washed with PBS and fixed with 4% paraformaldehyde. A dual fluorescent staining solution was prepared containing 100 μg/mL AO and 100 μg/mL PI. Cellular and nuclear morphologies were analyzed using a Nikon Eclipse E200 Epifluorescence microscope.

### Clonogenicity method

MDA-MB-231 cells were grown until an 80-90% confluent monolayer was reached and treated with different concentrations of the copper(II) complexes for 24 h. Subsequently, the cells were detached using TrypLE™, diluted, and seeded at approximately 1000 cells per well in 12-well plates. Cells were incubated for approximately 7 days, or until untreated control wells developed colonies containing at least 20 cells. Colonies were washed with PBS and stained with 0.5% crystal violet for 30 min at room temperature. Colonies were counted using an Olympus BX51 inverted microscope.

### Wound healing assay

MDA-MB-231 cells were cultured in 12-well plates in DMEM/F-12 medium supplemented with 10% FBS to achieve a 100% confluent monolayer. A linear scratch wound was made across the monolayer, washed with PBS, and treated for 24 h with 0.5 and 1.0 μM copper(II) complexes. The wells were washed with PBS and stained with Giemsa. We acquired images of the wounded area using an Olympus BX51 inverted microscope equipped with a digital camera. Cell migration was analyzed using ImageJ Software. The migration percentage was calculated using the following formula:

Wounded area= (final wound area (Tf) – initial wound area (T0) / initial wound area (T0)

### pH measurement and ammonium pulse

The NHE1 activity was assessed by evaluating pHi recovery following acute intracellular acidification induced by an ammonium chloride (NH₄Cl) prepulse. Intracellular pH (pHi) was measured by BCECF-AM (2′,7′-Bis-(2-Carboxyethyl)-5-(and-6)-Carboxyfluorescein, Acetoxymethyl Ester)-epifluorescence technique [28]. Briefly, MDA-MB-231 cells were seeded in 48-well plates and incubated overnight. Copper(II) complexes were added at the IC_50_ for 24 h. After treatment, cells were incubated for 15 min at 37°C with 10 μM BCECF-AM dissolved in Krebs-Henseleit (KH) solution (in mM): 146.2 NaCl, 4.7 KCl, 1 CaCl_2_, 10 HEPES, 0.35 NaH_2_PO_4_, 1 MgSO_4_, and 10 glucose. Steady-state pH was measured for 10 min in KH solution, next transient exposure to 20 mM NH_4_Cl was made for 10 min, and finally, the recovery was recorded for 15 min in KH solution. BCECF-AM fluorescence was measured through dual excitation (440 and 485 nm) and emission at 520 nm in a Varioskan LUX spectrofluorometer plate reader (Thermofisher). The 495-to-440 nm fluorescence ratio was calculated, and, at the end of each experiment, the fluorescence ratio was converted to pH by calibration using the high K^+^-nigericin method [29].

### Gelatin Zymography assay

The Gelatin Zymography assay was performed according to the adapted method of [30] to assess metalloproteinase (MMP) activity. Cells were seeded in 12-well plates and treated with the copper(II) complexes at their respective IC₅₀ for 24 h. Subsequently, the culture medium containing the complexes was removed, and cells were washed before overnight incubation in serum-free culture medium at 37°C. The cell-conditioned media were then collected, mixed with non-reducing loading buffer without β-mercaptoethanol, and subjected to electrophoresis on a 10% acrylamide gel containing 0.1% gelatin. After electrophoresis, the gels were washed with 2.5% Triton X-100 to remove SDS and allow enzyme renaturation. The gels were incubated at 37°C for 48 h in a buffer (50 mM Tris, 200 mM NaCl, 5 mM CaCl_2_, Triton X-100, pH 7.4) to allow gelatin degradation. Gels were stained with 0.5% Coomassie Brilliant Blue and destained with a methanol/acetic acid/water destaining solution (30:10:60, v/v/v). Gelatinase activity was quantified using Image J software and normalized to total protein.

### Western blotting

To determine the expression of metalloproteinase MMP-2, Bax, Bcl-2, NHE1, and GPER, cells were seeded in 6-well plates overnight and treated with the complexes for 24 h at their respective IC₅₀. Next, cells were lysed by adding cold RIPA buffer (KH_2_PO_4_ 30 mM, NaF 25 mM, EDTA 5 mM, sacarose 300 mM, Triton X-100 0.01 %, Igepal 1 %) with a protease inhibitor cocktail (Roche). 60 μg of protein (measured by the Bradford method [26] were loaded onto an 8 % SDS-polyacrylamide separating gel and transferred to polyvinylidene difluoride membranes (PVDF). Each membrane was incubated with specific primary antibodies against Bax, Bcl-2, NHE-1, MMP-2, GPER, and Na⁺/K⁺-ATPase (internal control) (1:1000; Santa Cruz Biotechnology). Next, membranes were incubated with anti-mouse secondary antibodies (1:10000; Cell Signaling). Immunoreactivity was visualized by a peroxidase-based chemiluminescence detection kit (Immobilon Western Millipore) using a Chemidoc Imaging System. The signal intensity of the bands in the immunoblots was quantified by densitometry using Image J software (NIH, USA).

### Statistical analysis

Results are expressed as the mean of independent experiments and represented as mean ± standard error of the mean (SEM). The total number of repeats (n) is specified in the legends of the figures. The Tukey test (two-way ANOVA) was employed to compare means in all the experiments performed.

## Results

### MDA-MD-231 cell viability

Cell viability of copper(II) complexes was studied using the MTT assay in the human breast cancer (MDA-MB-231) cell line, compared with Hacat (human keratinocyte) cells [14]. The results were compared with the reference metallodrug cisplatin. Our findings showed that the increase in methyl groups in the ligands (**2** and **3**) reduced MDA-MB-231 cell viability at lower concentrations (0.25 μM) than the complex with no methyl groups (Fig.2).

**Figure 2.**
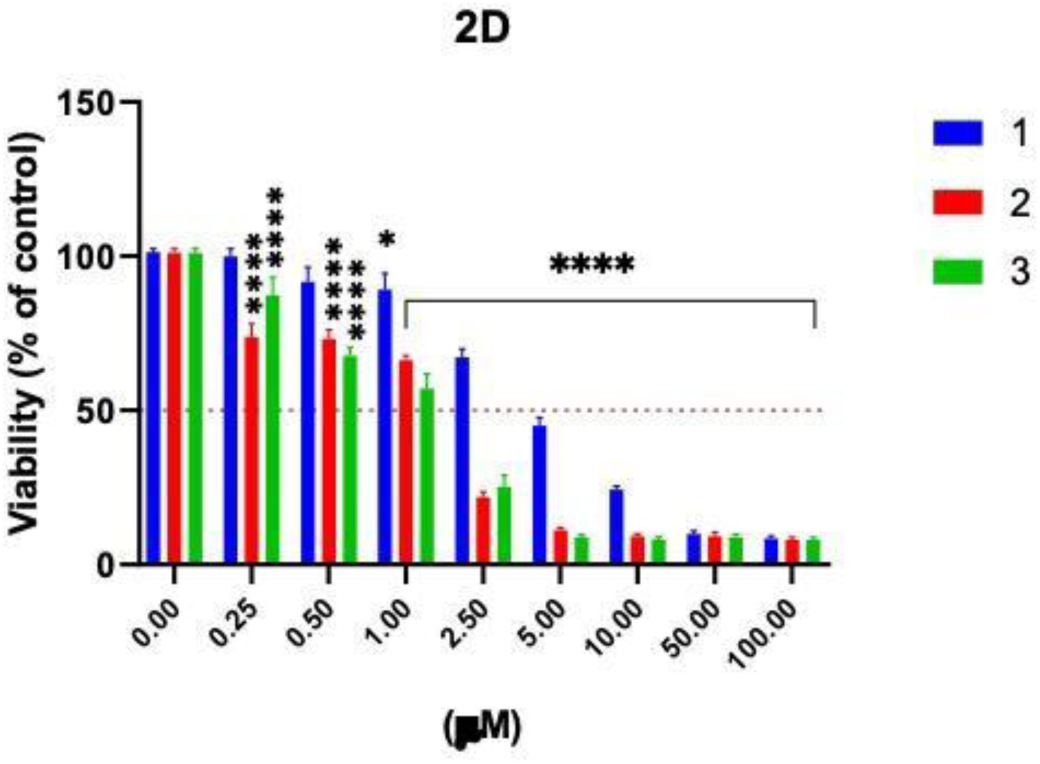
Cytotoxicity assay in breast cancer cells (MDA-MB-231) with different concentrations of the complexes at 37°C for 24 h, copper salt, and their ligands. Results are expressed as a percentage of the basal level and represent the mean ± standard error of the mean (SEM) (n=16). Asterisks represent a statistically significant difference in comparison with the basal level ****(p<0.0001).

The three complexes exhibited lower IC_50_ values than the corresponding ligand, the copper(II) ion, and cisplatin. This suggests that complexation is a key point in the antitumor properties. (Table 1).

**Table 1.** IC_50_ values of MDA-MB-231 and Hacat cells (24 h) for the three copper(II) complexes, a copper salt, phenanthroline, neocuproine, tetramethylphenantroline and cisplatin as positive control. **** (p<0.0001) and * (p<0.05). ^a^ Data previously reported by our group [14] ^b^ Selectivity index (SI) is IC50Hacat/IC50 MDA-MB-231. ^C^ [39]

| Complexes | $\text{IC}_{50}$ MDA-MB-231<br>(24 h) | $\text{IC}_{50}$ HaCaT<br>(24 h)<sup>a</sup> | SI<sup>b</sup> |
| --- | --- | --- | --- |
| **1** | $3.7 \pm 0.5$ | $5.0 \pm 0.7$ | 1.3 |
| **2** | $1.8 \pm 0.2$ | $1.8 \pm 0.2$ | 1.0 |
| **3** | $1.6 \pm 0.2$ | $2.4 \pm 0.4$ | 1.4 |
| **CuCl<sub>2</sub>** | > 100 (7) | - | - |
| **phen** | $48.5 \pm 1.6$ | $11.7 \pm 1.1$ | 0.2 |
| **neo** | $82.5 \pm 1.9$ | $21.7 \pm 3.3$ | 0.3 |
| **tmp** | $156.6 \pm 2.1$ | $9.3 \pm 2.4$ | 0.1 |
| **cisplatin** | $27.0 \pm 1.9^c$ | $24.4 \pm 6.9^c$ | 0.9 |

**Table 2.** Intracellular copper values in MDA-MB-231 (24 h) after treatment with IC_50_ of the three complexes and CuCl_2_ as control.

| Complexes | Cu(ng)/protein(mg) |
| --- | --- |
| **1** | 1257.88 |
| **2** | 2551.38 |
| **3** | 1883.29 |
| **CuCl<sub>2</sub>** | 718.67 |

Moreover, the copper(II) complexes exhibited moderate selectivity for tumor cells, sufficient to support further mechanistic studies, particularly given their high cytotoxic potency.

### Genotoxic effect

DNA damage induced by copper(II) complexes was studied using the alkaline comet assay, in which damaged DNA migrates out of the nucleus, forming a tail that can be quantified by image analysis. MDA-MB-231 cells were exposed to concentrations lower than the IC_50_ of all complexes (1.0 and 2.0 μM). Only 2 μM of **1** showed pronounced DNA damage (Fig. 3).

**Figure 3.**
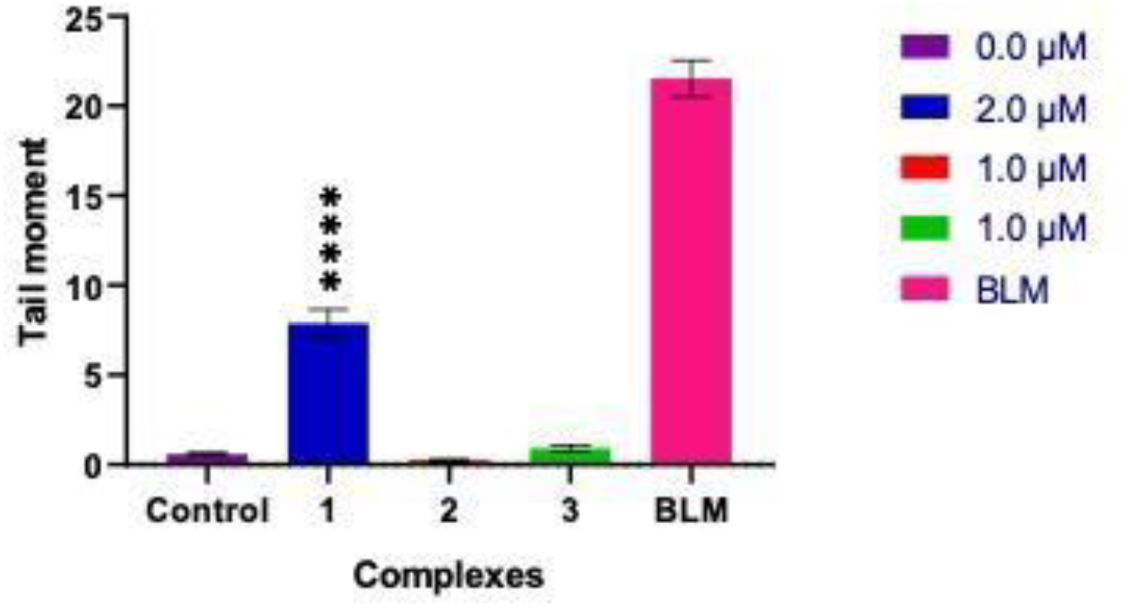
DNA damage (Tail moment) in MDA-MB-231 cells. Results are expressed as the mean ± SD (n=100). **** (p<0.0001).

### Oxidative stress induction

ROS production was evaluated using the probe DHR-123 to measure intracellular H_2_O_2_. The results showed that 0.5 and 1.0 μM of **2** and **3** induced oxidative stress, with DHR-123 oxidation increasing by 129 % for both complexes at 0.5 μM and by 121% and 112% at 1.0 μM for **2** and **3**, respectively. Complex **3** presented the highest ROS level at both concentrations (Fig. 4A).

**Figure 4.**
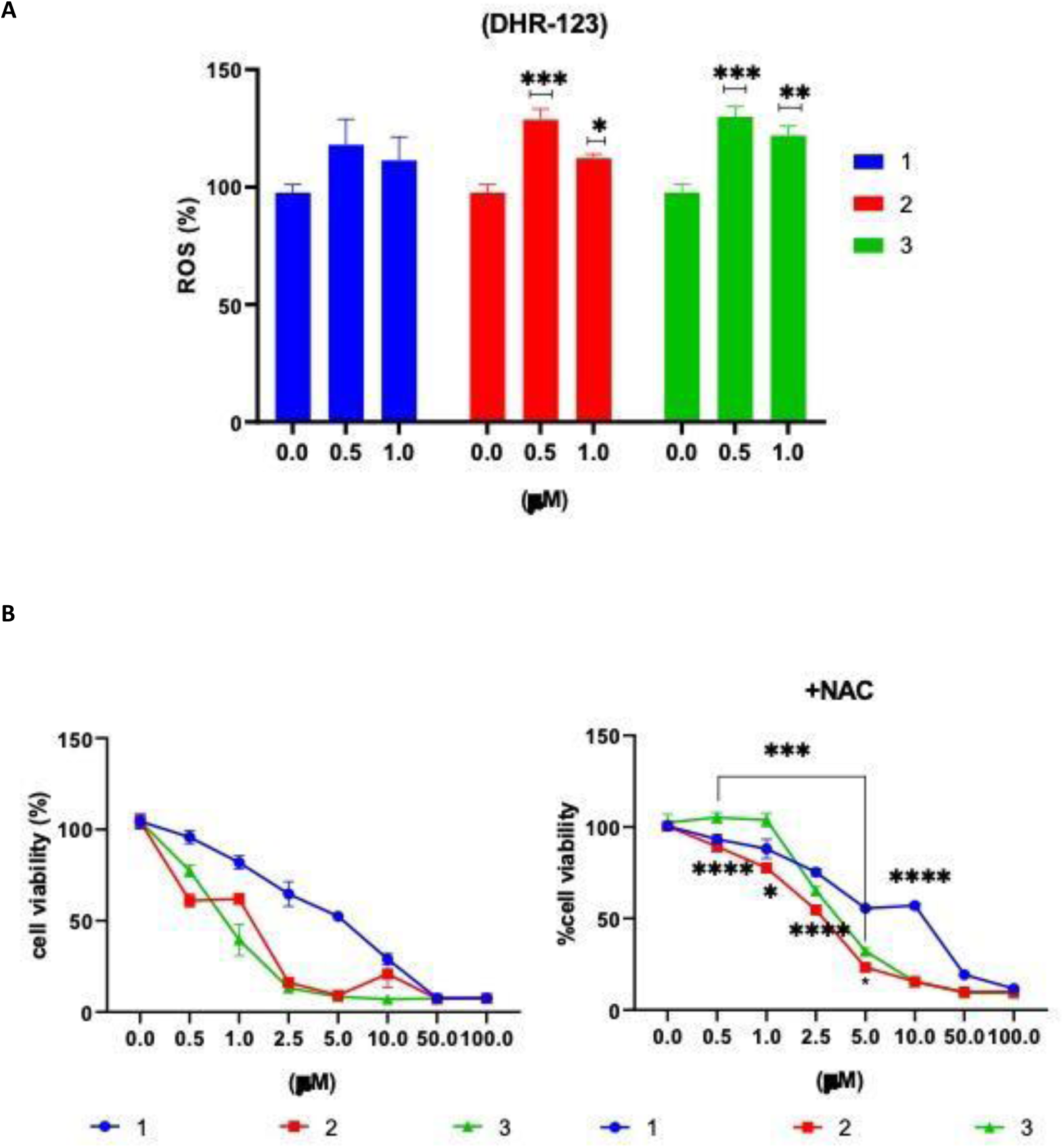
**A**. ROS level in the MDA-MB-231 cell line evaluated through the oxidation of dihydrorhodamine 123 (DHR-123) (n=3). **B**. Influence of the addition of N-acetyl cysteine in MDA-MB-231cells. Cell viability differed significantly among NAC-treated cells at different concentrations (n =3). **** (p<0.0001), *** (p<0.001) and * (p<0.05).

Exogenous antioxidant scavengers, such as NAC (N-Acetyl-L-Cysteine), recovered cell viability in **2** and **3**, suggesting that ROS production may contribute to the complexes’ cytotoxicity in this cell line (Fig. 4B).

In this context, enzymatic and antioxidant biomarkers were used to evaluate and confirm the presence of ROS, namely glutathione (GSH), superoxide dismutase (SOD), and catalase (CAT) (Fig. 5).

**Figure 5.**
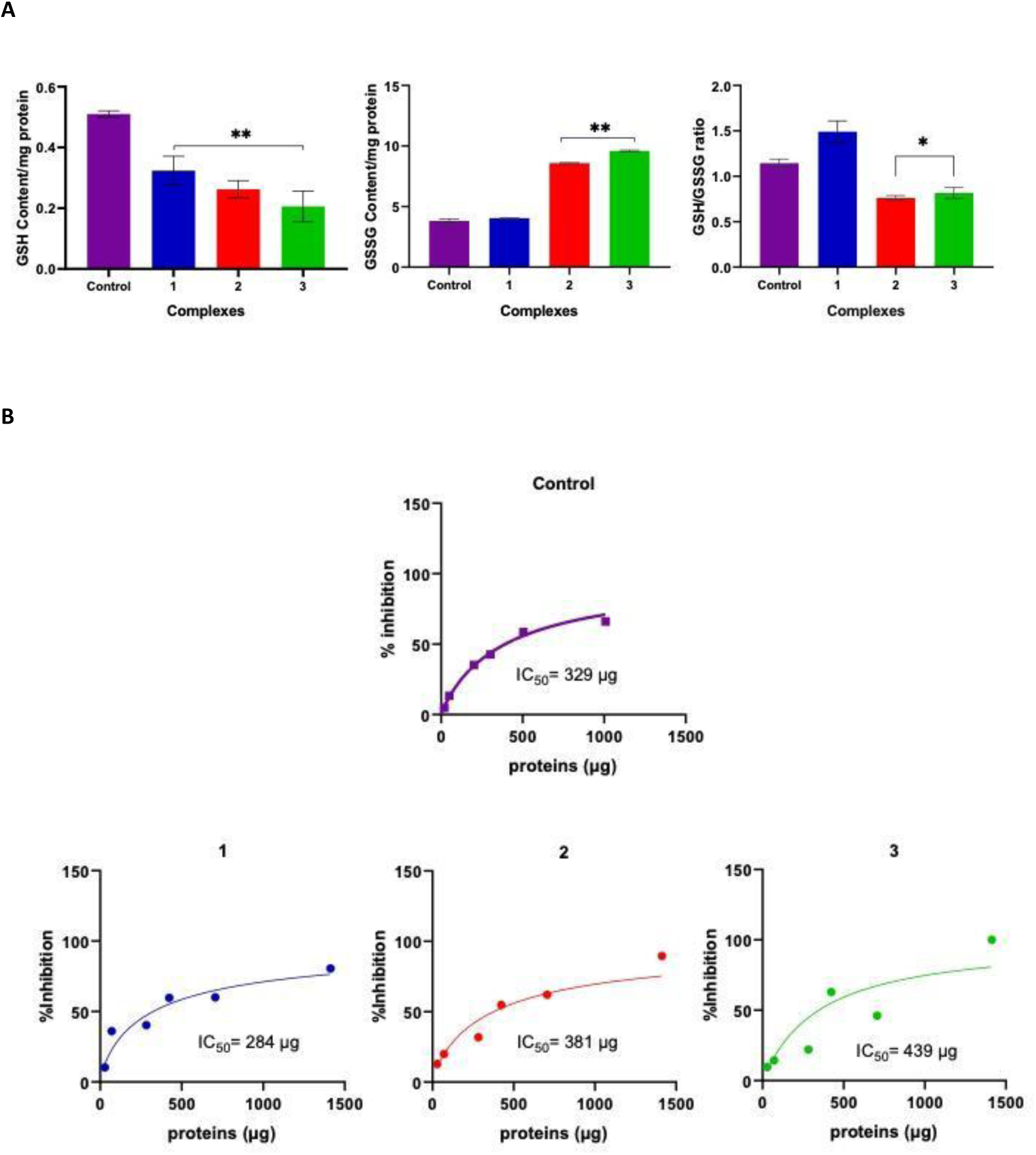
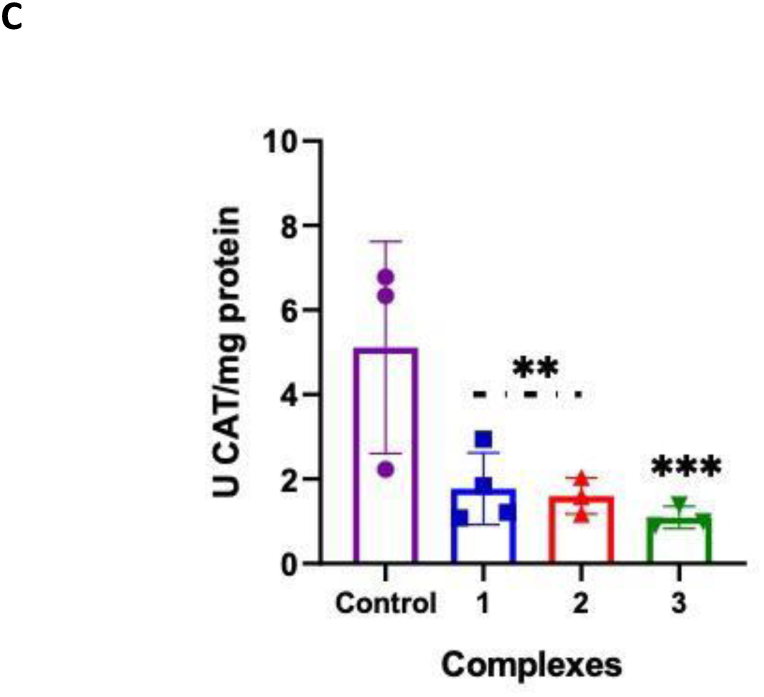
**A.** GSH level, GSSG level, and GSH/GSSG ratio **B**. IC_50_ value superoxide dismutase (SOD) and **C**. Catalase (CAT) activity in MDA-MB-231 cells. Results are expressed as the mean ± the standard error of the mean (SEM) (n=3), *** (p<0.001) ** (p<0.01) and * (p<0.05).

GSH is a multifunctional molecule that plays a key role in several cellular processes, mainly cell survival. When reactive oxygen species such as hydrogen peroxide are present, glutathione peroxidase metabolizes them, forming GSSH. Both GSH concentration and the GSH/GSSG molar ratio help maintain redox balance within the cell. We observed a decrease in GSH across all complexes; however, GSSH accumulation indicates active metabolism in response to ROS generated by **2** and **3**, as confirmed by a low GSH/GSSH ratio.

The SOD and catalase systems were inhibited by **2** and **3**. The ability of SOD to eliminate superoxide radicals was decreased by increasing the IC_50_ of SOD (the amount of sample needed to inhibit 50% of the signal produced by superoxide radicals) compared with the IC_50_ SOD control. These complexes decrease cellular antioxidant capacity, while **1** shows partial inhibition.

### Apoptosis induction

To establish the cell death mechanisms of the copper(II) complexes, an acridine orange (AO) and propidium iodide (PI) staining assay was performed. Green fluorescence in living cells, orange fluorescence during apoptosis, and red fluorescence in necrotic cells are the color patterns, since AO penetrates cells (intercalating with DNA), and PI is an intercalating agent in altered cell membranes.

All the complexes induced early apoptosis, and only **3** (highest number of methyl groups) exhibited high cytotoxicity, triggering both regulated apoptosis and membrane disruption (necrosis) at the IC_50_ (Fig. 6).

**Figure 6.**
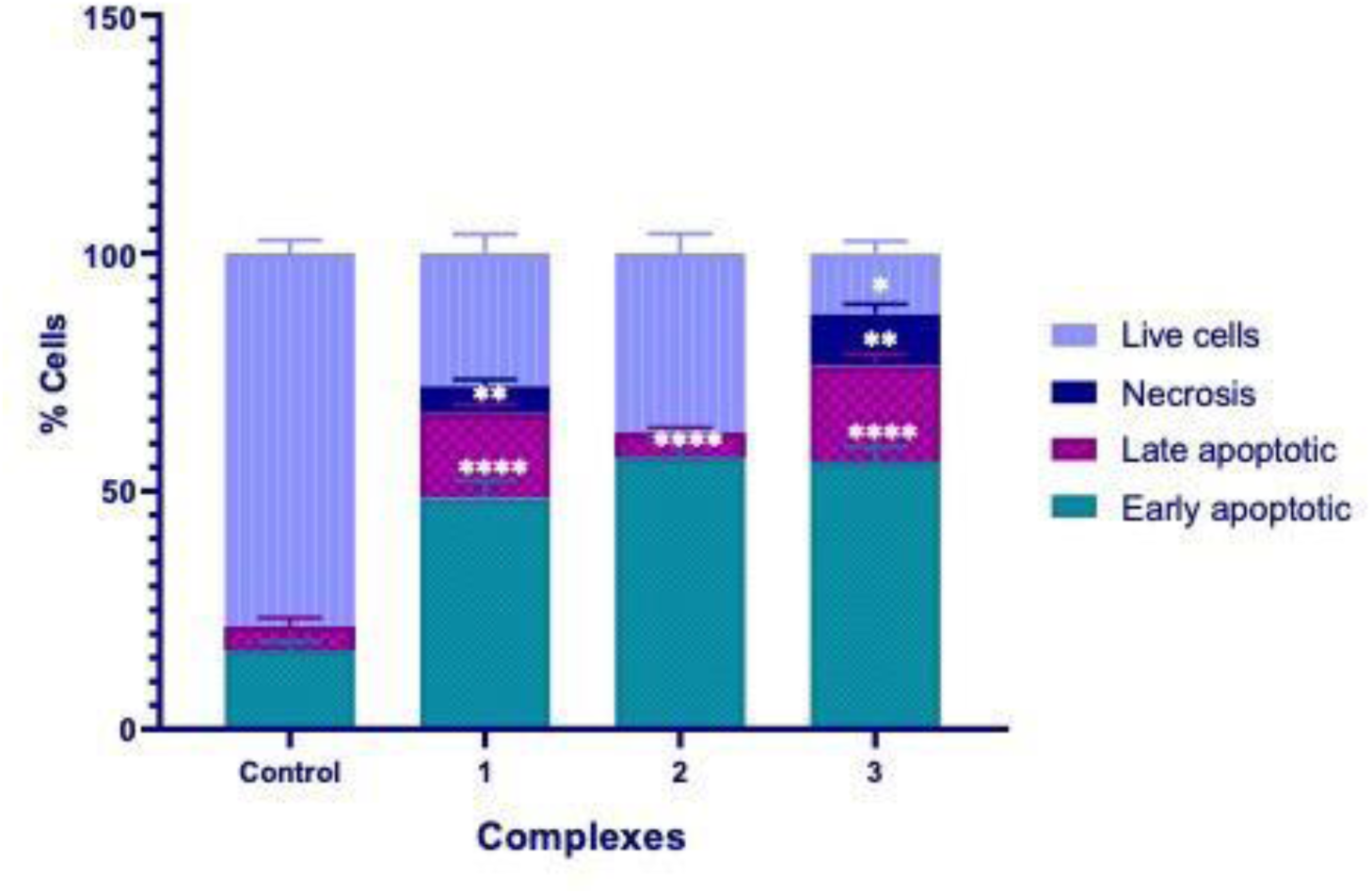
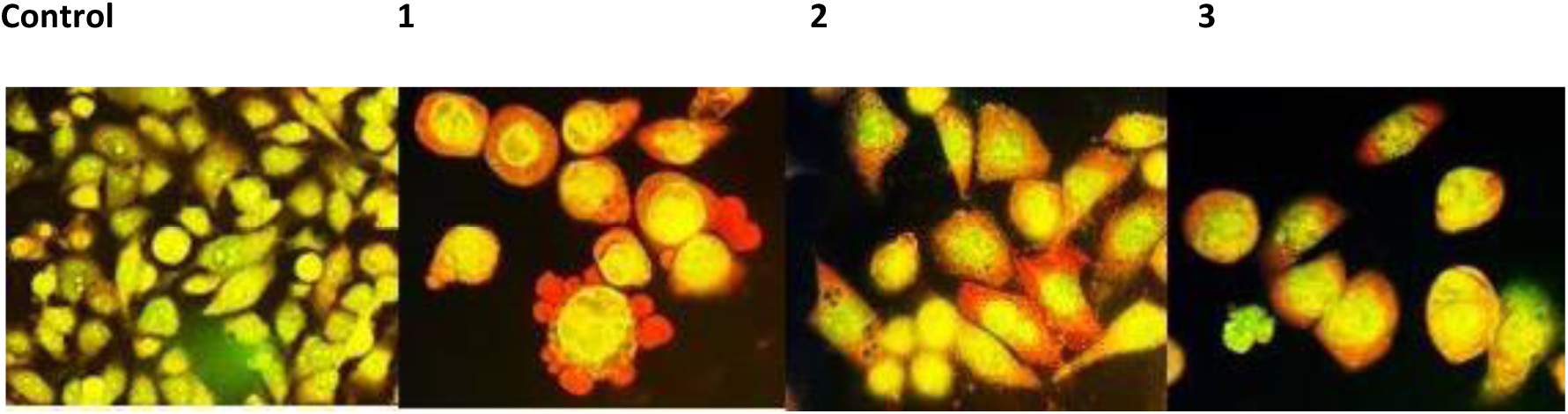
**A.** Cell death was analyzed by AO/PI stained cells. **B.** Cell morphology in MDA-MB-231 using acridine orange (AO) and propidium iodide (PI) to determine the percentage of cell apoptosis. **** (p<0.0001), ** (p<0.01) and * (p<0.05).

### Reduction of cell survival and migration

The clonogenic assay is a cell survival assay based on a single cell’s ability to grow into a colony. We observed that all complexes affected clonogenic capacity after a 24 h treatment; however, complexes **2** and **3** reduced the number of clones from 1.0 μM (100 and 47%, respectively), and **1** from 3 μM (Fig. 7). To analyze whether complexes reduce migration, we used a wound-healing assay. Our results showed that only **1** did not inhibit cell migration in the tumor cell line studied; **2** and **3** reduced cell migration from 1 and 0.5 μM (Fig. 7).

**Figure 7.**
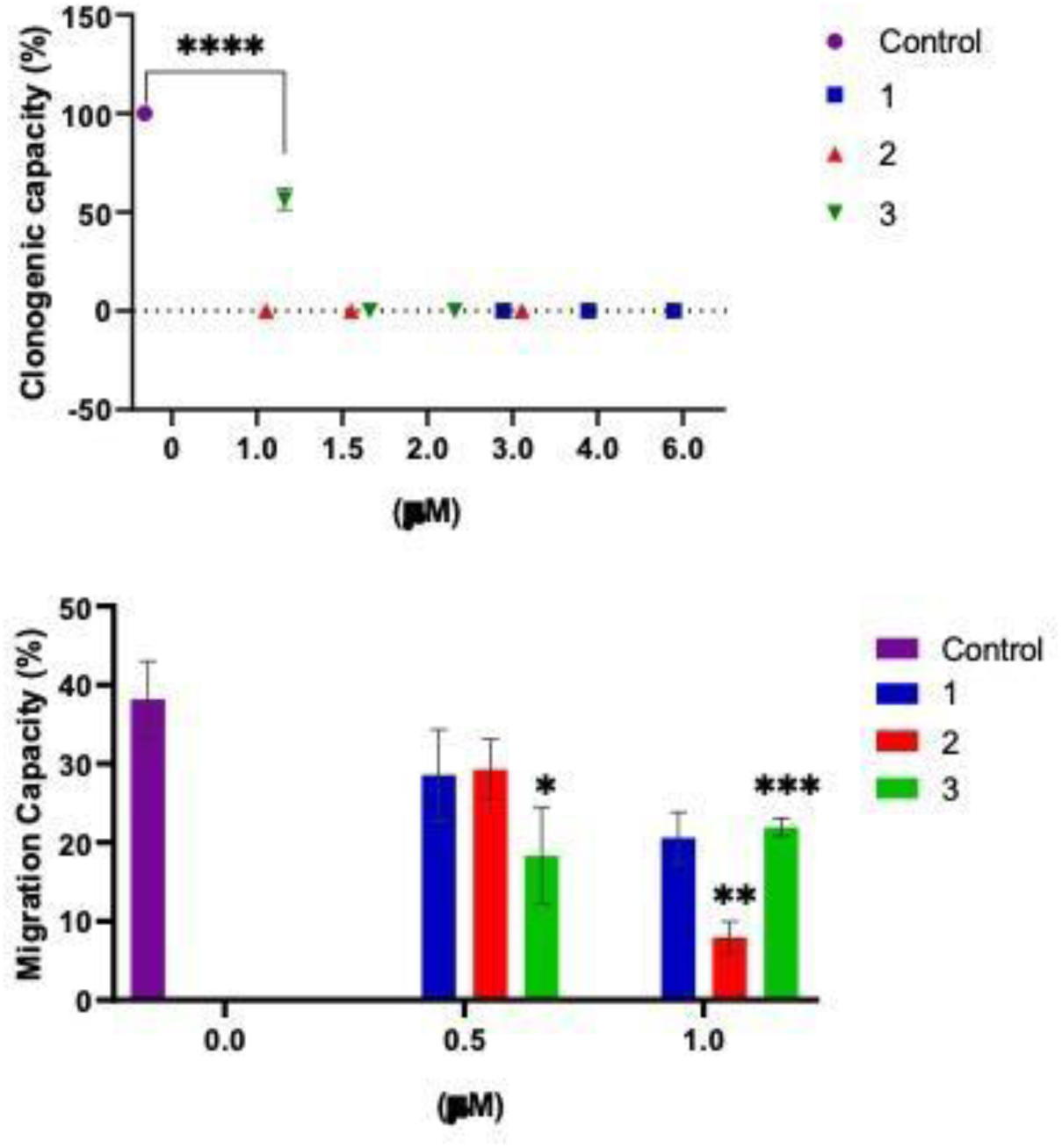
Clonogenic capacity and cell migration in breast cancer cells (MDA-MB-231) with different concentrations of the complexes at 37°C for 24 h. The results are expressed as the percentage of the basal level and represent the mean ± the standard error of the mean (SEM) (n=3) **** (p<0.0001) *** (p<0.001), ** (p<0.01) and * (p<0.05).

### Intracellular copper

Copper is vital for cell growth and development; however, slight alterations in its homeostasis can cause severe toxicity, activate apoptosis, and lead to cell death [40]. Intracellular copper concentration increased when tumor cells were exposed to the three complexes and rose threefold with **2** compared with the metal salt control.

### NHE-1 activity

NHE-1 is an important exchanger that plays a fundamental role in tumor growth, particularly in MDA-MB-231 cells. Our findings show that all the complexes examined inhibited this exchanger without increasing intracellular pH (pHi) at 5 minutes, compared with the control (Fig. 8). The effect of inhibition was more pronounced for **2** and **3**, leading to intracellular acidification. This result is consistent with computational studies in which this complex exhibited a higher binding free energy for this exchange (see below).

**Figure 8.**
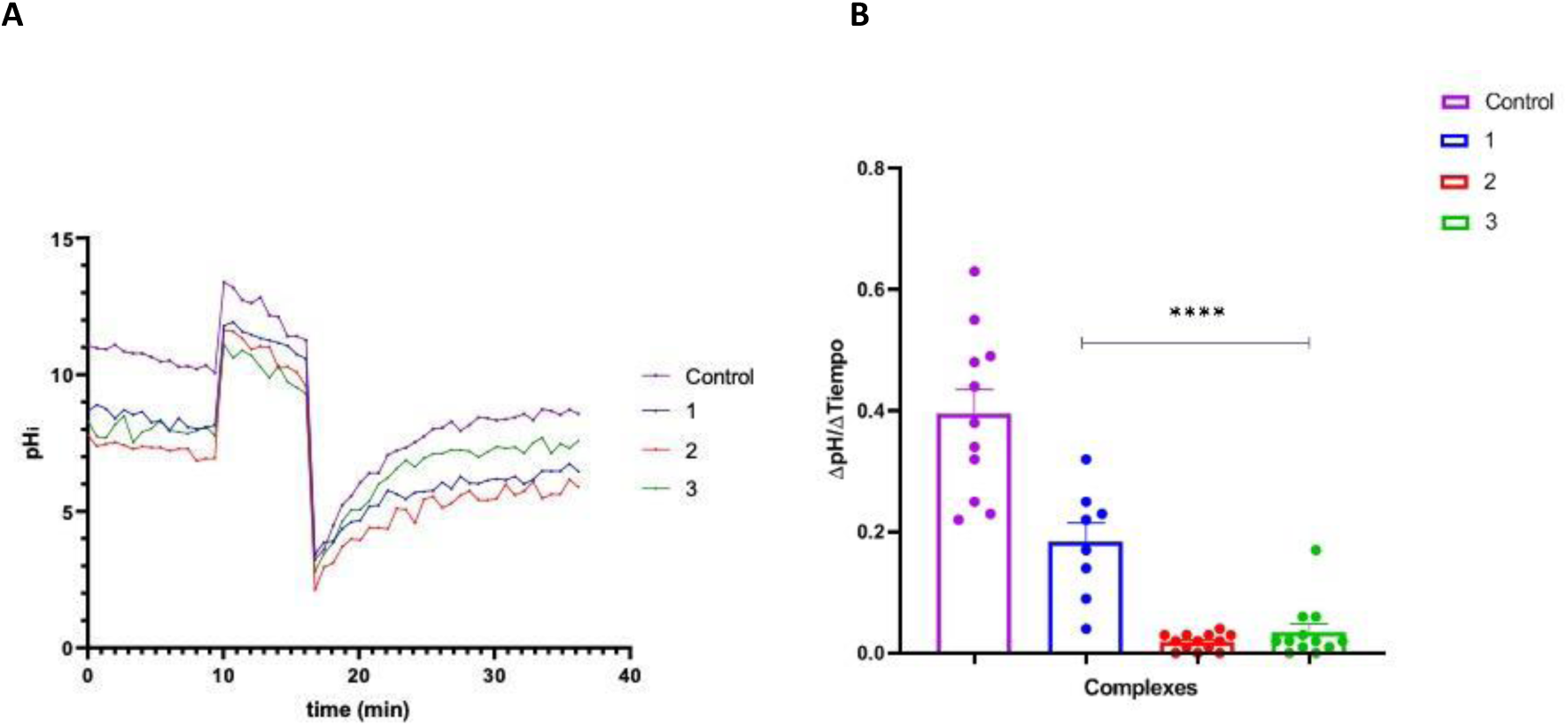
**A**. Representative traces of pHi recovery after the ammonium pulse (20 mM NH_4_Cl) in MDA-MB-231 treated with the complexes at the IC_50_ and perfused with HK buffer. **B**. Average proton efflux (JH) carried by NHE-1, calculated at different pHi values during the recovery from acidosis. Results are expressed as a percentage of the basal level and represent the mean ± standard error of the mean (SEM) (n=3) * (p<0.05).

### Effect of copper(II) complexes on GPER, Bax, BCL-2, NHE1, MMP-2 expression

The effect of the three complexes on Bax, Bcl-2, MMP-2, GPER, and NHE-1 expression was observed by Western blotting analysis (Fig.9). None of the complexes statistically inhibited MMP2. However, all the complexes inhibited GPER and NHE-1. Similarly, they inhibited apoptosis-regulating proteins, both Bax (a pro-apoptotic protein) and Bcl-2 (an anti-apoptotic protein), with the latter inhibited significantly. However, the tendency to exhibit a high Bax/Bcl-2 ratio indicates that **2** and **3** operate via a mechanism in which oxidative stress induced by these copper(II) complexes shifts the balance between pro- and anti-apoptotic proteins, favoring mitochondrial apoptosis (intrinsic pathway).

**Figure 9.**
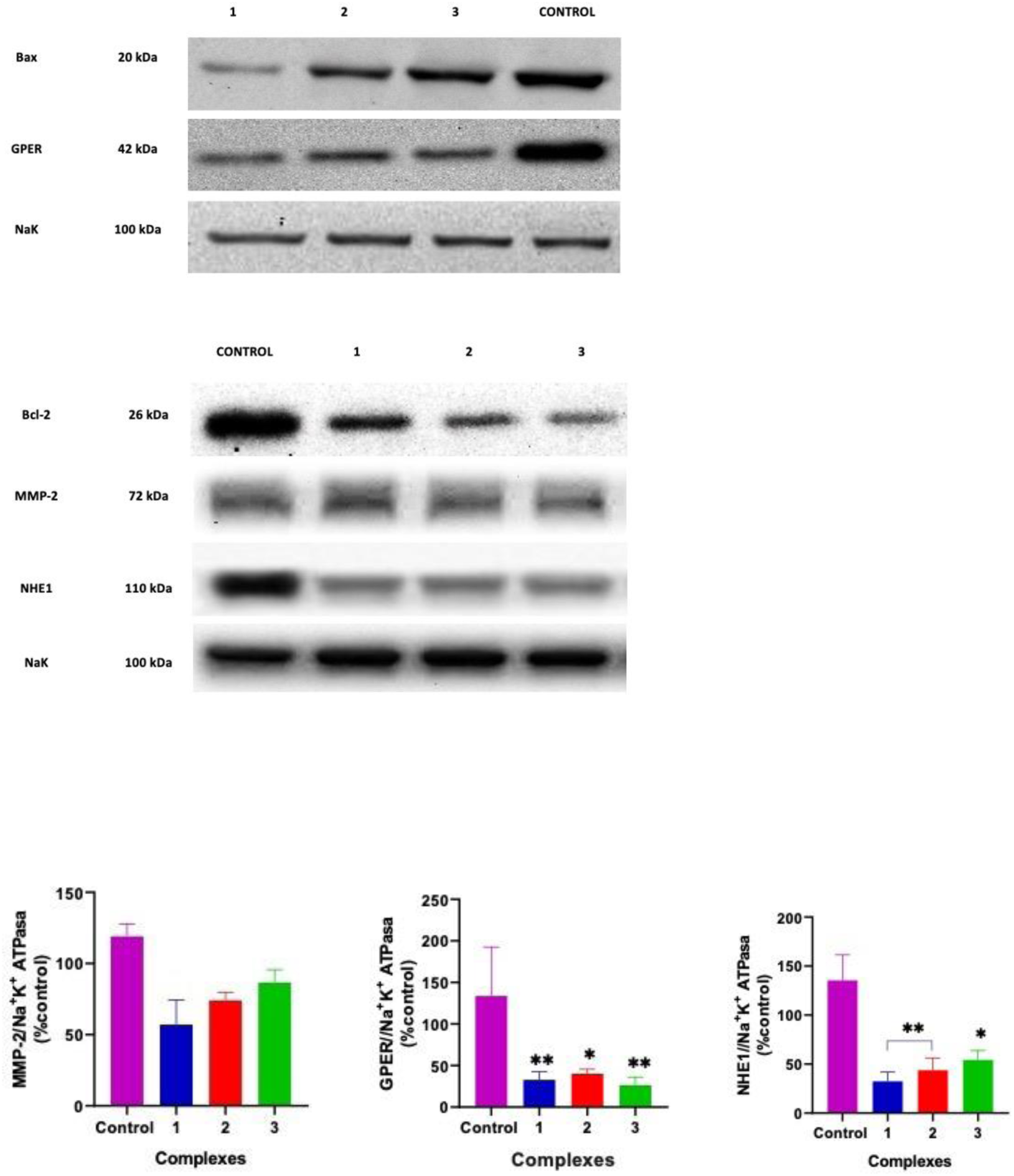
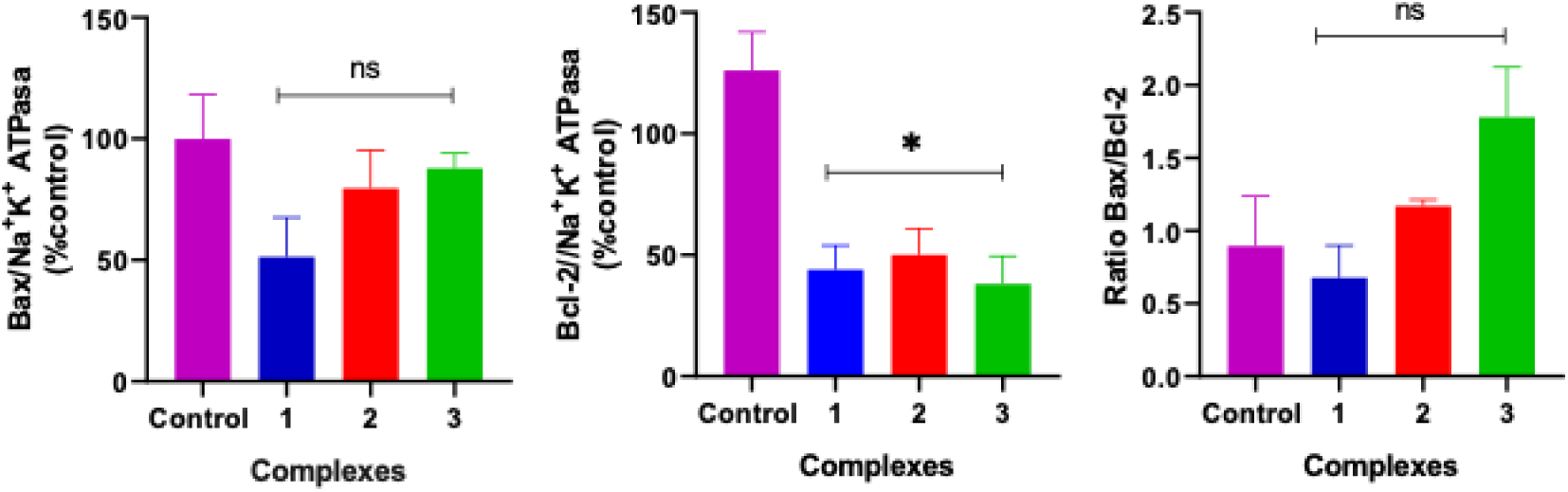
Western blot analysis of the three copper(II) complexes for BAX, Bcl-2, Ratio Bax/Bcl-2, MMP-2, GPER, and NHE1. Results are expressed as the mean ± standard error of the mean (SEM) (n = 3). Asterisks indicate significant differences with the control * (p < 0.05), ** (p < 0.01), *** (p < 0.001), **** (p < 0.0001).

### Effect of copper(II) complexes on MMP-9 and MMP-2 activity

Given the effects observed on cell migration, we evaluated MMP-2 and MMP-9 activity to further investigate the potential role of copper complexes in limiting cell migration and invasion. MMP-2 and MMP-9 are key gelatinases that degrade extracellular matrix components and have been associated with tumor cell migration and invasive behavior [41–43]. Our results indicate that the three complexes inhibited pro-MMP-9 and active MMP-9. However, neither affected pro-MMP-2 nor active-MMP-2 (Fig. 10).

**Figure 10.**
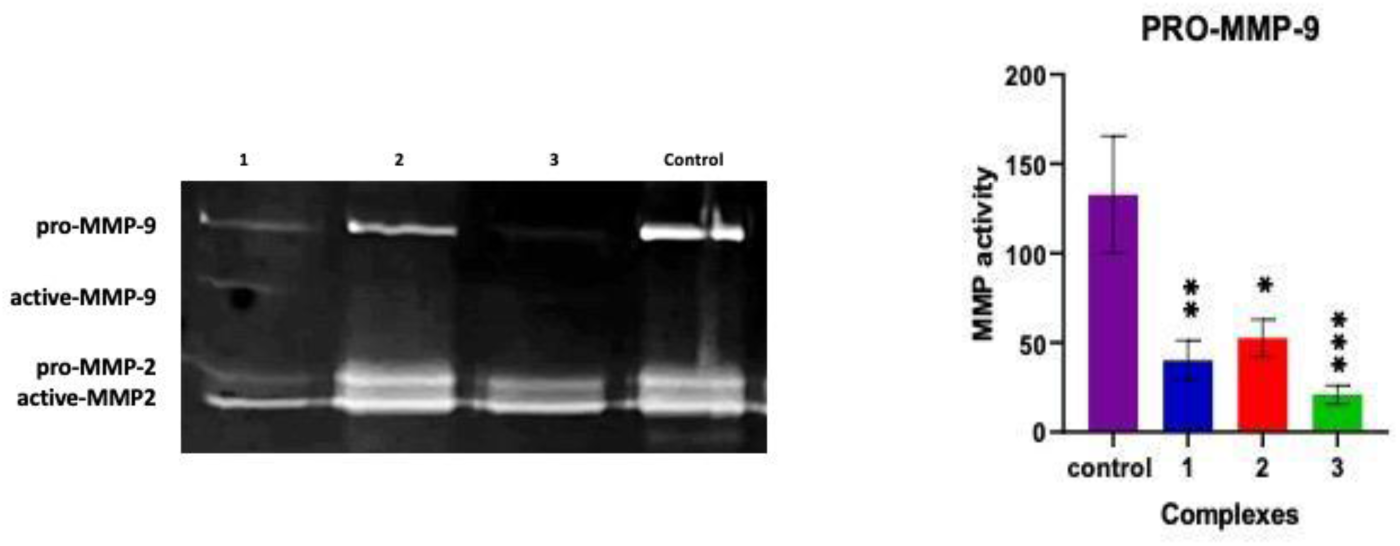
Pro-MMP-9 activity by zymography in MDA-MB-231 extracts after 24 h. Results are expressed as the mean ± standard error of the mean (SEM) (n = 3). Asterisks indicate significant differences with the control: * (p < 0.05), ** (p < 0.01), *** (p < 0.001).

## Discussion

### Copper coordination as a key determinant of antitumor activity

The development of copper-based anticancer agents has attracted increasing attention as an alternative to platinum-derived drugs because of their multiple mechanisms of action, lower susceptibility to drug resistance, and ability to alter copper homeostasis in many tumors [44]. In this context, phenanthroline-containing copper(II) complexes have emerged as particularly attractive candidates because they combine the redox activity of copper with the DNA-binding and lipophilic properties of diimine ligands [4,45].

One of the most relevant findings is that coordination of copper(II) with phenanthroline derivatives dramatically enhances cytotoxic activity compared with both the free ligands and the copper salt alone. Whereas Cu²⁺ ions and the unconjugated ligands exhibited only limited cytotoxicity, all three copper(II) complexes showed low micromolar IC₅₀ values and were more potent than cisplatin under identical experimental conditions. These results clearly demonstrate that complex formation is essential for the observed biological activity. Consistent with our findings, Mohindru et al. reported that both the cytotoxicity and cellular uptake of neocuproine strongly depend on copper coordination, highlighting the critical role of metal complexation [46]. Smolková et al. showed cytotoxicity across different transition metals, and the reported copper complex was more effective against MDA-MB-468 (TNBC) and HCT116 (colorectal cancer) cells [47].

Moreover, methyl groups play an important role in their antitumor properties. The IC_50_ value decreases as the number of methyl groups in the molecule increases. Complexes with two methyl groups (**2**) and four methyl groups (**3**) inhibited cell viability below 1.5 µM, indicating a clear structure–activity relationship. Such effects have been described previously for methylated phenanthrolines, particularly neocuproine and tetramethylphenanthroline derivatives, which frequently exhibit enhanced biological activity compared with unsubstituted phenanthroline. This effect may be due, at least in part, to increased lipophilicity conferred by methyl substitution, which facilitates cellular uptake and improves intracellular copper delivery [48].

Selectivity also deserves consideration. Although the three complexes exhibited only modest SI values, these were consistently higher than those of the corresponding free ligands or cisplatin, indicating that copper coordination not only enhances cytotoxic potency but also improves the therapeutic window, *i.e.*, the balance between antitumor efficacy and toxicity toward non-malignant cells.

Thus, these results show that subtle structural modifications of the phenanthroline scaffold and copper(II) complexation markedly influence biological activity.

Our findings suggest that cytotoxic potency cannot be explained solely by DNA damage. The genotoxic effects of copper complexes investigated using the Comet assay demonstrated that only **1** induced significant DNA damage. Among compounds exhibiting DNA cleavage activity (nuclease-like), the [Cu(phen)₂]²⁺ is the most extensively studied. Sigman and co-workers first described it, demonstrating the reduction to [Cu(phen)₂]⁺, which binds to the minor groove of DNA. The reduced copper complex reacts with molecular oxygen to generate oxidizing species that promote DNA strand scission by oxidizing the deoxyribose backbone[49]. Previous studies suggest that adding methyl groups to a phenanthroline ligand often impairs its ability to bind DNA, primarily due to steric interference. The bulky methyl groups disrupt the planarity and prevent the ligand from intercalating between DNA base pairs, which may contribute to the reduced genotoxicity observed for the methylated derivatives [50,51]

These results are particularly relevant because they support the emerging concept that modern metallodrugs should not necessarily function as classical DNA-targeting agents, as with cisplatin, but rather act by simultaneously modulating multiple signaling pathways involved in tumor progression [52].

### Copper accumulation, oxidative stress, and apoptosis constitute the central mechanism underlying the cytotoxic activity of the copper(II) complexes

Intracellular copper homeostasis is tightly regulated. Cancer cells are particularly susceptible to disturbances in copper metabolism because they frequently exhibit increased copper uptake and a greater dependence on copper-dependent signaling pathways than their non-malignant counterparts. However, exceeding a critical threshold triggers cuproptosis (copper-induced cell death) and irreversible oxidative damage [53]. Consequently, copper-based complexes can exploit this metabolic vulnerability by selectively increasing intracellular copper levels beyond tumor cells’ buffering capacity [44]. The present results demonstrate that the three copper complexes exhibited markedly different intracellular copper accumulation, with **2** producing nearly a three-fold increase compared with CuCl₂. This observation is particularly significant because it indicates that the biological activity of these compounds depends not only on their intrinsic chemical reactivity but also on their ability to efficiently deliver copper into the cell, a capacity that may be related to lipophilicity [54,55].

Consistent with these findings, several phenanthroline-containing copper complexes showed biological activity that correlates more closely with intracellular copper accumulation than with extracellular drug concentration [45,56]. The increase in intracellular copper was accompanied by a marked elevation in ROS, statistically significant for **2** and **3**, supporting the hypothesis that oxidative stress is a major determinant of their cytotoxic activity. The protective effect of N-acetylcysteine further supports this mechanism. Restoration of cell viability after antioxidant treatment strongly suggests that ROS generation is not merely a secondary consequence of cell death but rather an essential event in the cytotoxic mechanism. A comparable pattern has been reported for numerous redox-active copper complexes, in which antioxidant supplementation significantly attenuates cytotoxicity by limiting oxidative damage [57]. Additional support for oxidative stress-mediated toxicity comes from marked alterations in the endogenous antioxidant defense system. Glutathione is the principal intracellular redox buffer, and a decrease in reduced glutathione, together with the accumulation of oxidized glutathione, indicates that cellular antioxidant capacity becomes progressively overwhelmed following exposure to the copper complexes [58]. Consequently, the reduced GSH/GSSG ratio observed for **2** and **3** reflects a shift toward a pro-oxidant intracellular environment. Simultaneously, inhibition of superoxide dismutase and catalase further compromises cells’ ability to detoxify superoxide radicals and hydrogen peroxide, thereby amplifying oxidative injury [44]. Likewise, in our previous studies, we found that the cytotoxic effect of several copper complexes was associated with oxidative stress induction across different cancer cell lines, suggesting that ROS generation is a common feature contributing to the antitumor activity of copper(II) compounds [59–61]. In contrast, in MCF-7 cells, only the tmp-containing complex induced a significant increase in DHE fluorescence, indicating enhanced superoxide production, whereas the other two copper complexes did not alter DHE levels. Moreover, none of the complexes increased DHR123 oxidation, and NAC failed to restore cell viability. These findings suggest that in this cell line, ROS generation, when present, represents a secondary event rather than a key mechanism underlying the cytotoxic effects of the copper complexes [14]. Given that triple-negative breast cancer cells typically exhibit higher basal ROS levels and greater dependence on antioxidant defenses, they may be closer to a critical redox threshold, making them more susceptible to the additional oxidative burden imposed by redox-active copper complexes [62].

Interestingly, although both **2** and **3** induced oxidative stress, their downstream biological responses differed. Although both complexes activated apoptotic pathways, **3** additionally promoted membrane disruption and necrotic features. This finding suggests that increasing methyl substitution not only enhances oxidative stress but may also influence the balance between regulated apoptosis and secondary necrotic events [63]. Consequently, ligand structure appears to determine not only the magnitude of oxidative stress but also the cellular decision between different modes of cell death.

### Copper complexes impair long-term survival and suppress the invasive phenotype of breast cancer cells

The clonogenic assay is considered the gold standard for evaluating long-term cell survival because it measures a single cell’s ability to proliferate indefinitely and form a macroscopic colony after exposure to a cytotoxic agent [64]. While apoptosis induction is a desirable feature of antitumor agents, their ability to suppress long-term survival and metastasis is also relevant. In this study, the copper complexes markedly reduced clonogenic capacity, indicating that surviving cells lost their ability to proliferate indefinitely and exerted an antitumor effect beyond the acute cytotoxic response.

The reduction in cell migration further supports the potential of **2** and **3** to interfere with mechanisms associated with tumor progression. Cell migration is a highly coordinated process that depends on cytoskeletal remodeling, extracellular matrix degradation, and the establishment of intracellular pH gradients [65]. One of the main regulators of this process is NHE1, whose increased activity is a hallmark of many aggressive tumors. By extruding protons, NHE1 promotes intracellular alkalinization while acidifying the pericellular environment, conditions that favor activation of extracellular proteases [66]. Consistent with this concept, inhibition of NHE1 activity observed after treatment with **2** and **3** provides a plausible mechanistic link between NHE1 inhibition and the reduced migratory capacity observed after treatment.

Migration inhibition was accompanied by a decrease in pro-MMP-9 activity but did not affect pro-MMP-2 or active-MMP-2. These two gelatinases play a critical role in extracellular matrix degradation and cell invasion [67], supporting the relevance of reduced MMP-9 activity to the impaired cell migration observed in our study.

Interestingly, MMP-2 activity remained unaffected. This differential effect suggests that the compounds may preferentially interfere with mechanisms regulating MMP-9 activity rather than broadly suppressing gelatinase activity. Moreover, MMP-2 and MMP-9, despite sharing similar substrate specificities, may exert distinct functional roles in cell–matrix interactions and cell migration [68]. Studies in MDA-MB-231 cells have shown that MMP-2 and MMP-9 contribute differently to the invasive phenotype. Silencing MMP-2 reduced cell invasion by about 50%, whereas MMP-9 knockdown reduced it by nearly 90%, highlighting MMP-9’s particularly relevant role in tumor cell motility and invasion [69].

These findings align with previous studies showing that copper complexes reduce MMP activity through multiple mechanisms, including modulation of intracellular pH homeostasis via NHE1 inhibition and, in some cases, attenuation of ROS-dependent signaling pathways such as MAPK and NF-κB [60,70]. Western blot analysis further supports these functional findings. The increase in the Bax/Bcl-2 ratio is consistent with activation of the intrinsic apoptotic pathway. Moreover, reduced NHE1 expression correlates with reduced functional activity, suggesting that the complexes regulate this protein not only at the activity level but also at the expression level. Interestingly, modulation of GPER expression may also contribute to the observed biological effects. GPER has been implicated in breast cancer cell proliferation, migration, and resistance to therapy through the activation of ERK1/2, PI3K/Akt, and EGFR signaling pathways; thus, the reduction in GPER expression observed after treatment with the copper complexes may represent an additional molecular alteration associated with their antitumor activity. Its downregulation may help attenuate the aggressive phenotype observed in treated breast cancer cells [71].

Taken together, the three complexes exhibit distinct but interconnected mechanisms of antitumor action. **1** containing unsubstituted 1,10-phenanthroline, appears to rely more strongly on DNA damage-associated cytotoxicity. In contrast, methylated complexes **2** and **3** display a predominant redox-dependent mechanism accompanied by a more pronounced NHE1 inhibition and a selective reduction in migratory capacity. **2** is distinguished by efficient intracellular copper accumulation, whereas **3** combines strong oxidative stress with the most pronounced membrane-disruptive effects.

These findings indicate that methyl substitution does not simply increase cytotoxic potency but reshapes the relative contribution of DNA damage, redox imbalance, pH homeostasis, and migratory pathways to the overall antitumor response.

### Conclusion

Overall, our results reveal a clear structure–activity relationship, showing that ligand architecture modulates the antitumor properties of the copper complexes. Despite their comparable cytotoxic potency and selectivity, the compounds exhibited distinct DNA damage profiles, suggesting that subtle structural modifications influence their biological behavior.

Our findings support a mechanistic model in which efficient intracellular copper delivery initiates excessive ROS production, depletes antioxidant defenses, and activates apoptotic pathways. Moreover, our results show that copper(II) complexes suppress NHE1 function and reduce metalloproteinase activity, which may be central to subsequent inhibition of clonogenic potential and impairment of metastatic behavior.

This multitarget mechanism may be particularly advantageous for limiting tumor progression and metastatic dissemination, supporting these compounds as promising anticancer agents.

## Acknowledgments

This work was supported by UNLP (X1045, SX006, 11/X958), CONICET (PIP 0235), and ANPCyT (PICT 2021-00090, PICT 2021 − 00338) from Argentina.

